# Does Sleep Protect Memories Against Interference? A Failure to Replicate

**DOI:** 10.1101/705798

**Authors:** Carrie Bailes, Mary Caldwell, Erin J. Wamsley, Matthew A. Tucker

## Abstract

Across a broad spectrum of memory tasks, retention is superior following a night of sleep compared to a day of wake. However, this result alone does not clarify whether sleep merely slows the forgetting that would otherwise occur as a result of information processing during wakefulness, or whether sleep actually consolidates memories, protecting them from subsequent retroactive interference. Two influential studies (Ellenbogen, et al., 2006, 2009) suggested that sleep protects memories against the subsequent retroactive interference that occurs when participants learn new yet overlapping information (interference learning). In these studies, interference was much less detrimental to memory following a night of sleep compared to a day of wakefulness, a finding that provided strong evidence that sleep supports this important aspect of memory consolidation. In the current well-powered replication study, we repeated the protocol of Ellenbogen, et al. (2009) and, additionally, we examined the impact of intrinsic motivation on performance in sleep and wake participants. We were unable to replicate the finding that sleep protects memories against retroactive interference, with the detrimental effects of interference learning being essentially the same in wake and sleep participants. We also found that while intrinsic motivation benefitted task acquisition it was not a modulator of sleep-wake differences in memory processing. These finding of this replication study draw into question the claim that sleep protects memories against the effects of retroactive interference, and moreover, they highlight the importance of replicating key findings in the study of sleep’s impact on memory processing before drawing strong conclusions that drive the direction of future research.

## Introduction

Retention of newly learned declarative memories (e.g., fact-based information) improves following an interval filled with sleep as opposed to wakefulness [1–3]. However, there is considerable debate about how to characterize the role of sleep for memory processing, and specifically, whether sleep benefits memory primarily due to *active* or *passive* mechanisms [4].

A key feature of memory “consolidation”, as classically defined, is the development of increasing resistance to interference over time [5,6]. If sleep serves to stabilize newly acquired memories in this way, then information should not just be better remembered following sleep (compared to wake), but should also be more resistant to the interfering effects of new learning that occur after sleep has had a chance to consolidate these memories. It may be, however, that sleep does not stabilize memories, but merely slows the process of forgetting that would occur more rapid during wakefulness when the brain is more active and engaged in information processing. In this case, retention of information would still be superior to wake following sleep (without interference), but sleep would not protect the memory against interference (e.g., learning information that is similar to the originally-learned information), leading to a similar rate of subsequent forgetting regardless of whether sleep or wake preceded the interference [4].

The current replication is based on two influential studies that supported the hypothesis that sleep protects memory from subsequent interference, reporting that interference was less disruptive to memory when sleep, as opposed to wake, preceded interference learning [7,8].

### Sleep and Declarative Memory Processing

Although the term “consolidation” today has multiple meanings, memory consolidation was originally defined as the process by which an initial labile memory trace becomes increasingly resistant to retroactive interference over time [5,9]. According to this view, if memories are consolidated during sleep, then interfering information should have less of a negative impact on post-sleep memory retention, compared to a much stronger negative effect on post-wake retention.

From a neurobiological perspective, a number of candidate aspects of sleep have been suggested to play an active role in memory consolidation. For example, studies have observed correlations between time spent in stage 2 sleep or slow wave sleep (SWS) and declarative memory retention [10–12], suggesting that sleep stage-specific neural activity during these particular stages contributes to memory consolidation. Sleep-specific oscillations, such as sleep spindles [13,14] and other neuronal oscillations that occur during sleep, such as hippocampal ripples [15], have also been correlated with memory processing. While these studies are compelling, it should be noted that experimental manipulations of these aspects of sleep are still limited in number, and alternative hypotheses exist. Indeed, some suggest that sleep, like other physiological states (e.g., those induced with alcohol or benzodiazepines, for example), allows consolidation to occur simply because new learning is dramatically reduced during these times [16]. This line of reasoning suggests that sleep may be one of many possible states during which memory consolidation can occur, though it may be the most opportune time [17]. Thus, even if active processes of memory consolidation are occurring during sleep, this may not necessarily be due to specific properties of sleep or sleep-specific neuronal events.

From a behavioral perspective, a handful of studies have provided evidence that sleep may play an important role in stabilizing memories, making them more resistant to subsequent interference that occurs after sleep initially processed the memories [7,8,18]. This evidence that sleep protects memories against interference has been argued to indicate that *active* processes of consolidation must be occurring during sleep. Employing a classic interference learning protocol developed by Barnes and Underwood [19], two studies had participants learn a list of word pairs (A-B pairs) in the evening (prior to a night of sleep) or the morning (prior to a day of wake) [7,8]. When participants returned 12 hours later to be retested on some of the word pairs, those who had slept retained more word pairs that those who remained awake. Participants then underwent interference training for a subset of the originally-learned word pairs, followed by a test of the original B associates and the newly learned C associates. Interference training lead to a dramatic forgetting of the originally learned words in the wake participants, compared to those who slept.

However, subsequent research has failed to detect a similar effect of sleep on the susceptibility of new memories to interference. In one study using the same experimental protocol and paired associates task, the authors failed to see an effect of sleep on either recall of word pairs subject to interference learning [18]. However, participants in that study were required to answer correctly each word pair 3 times during training, as opposed to 1-2 times correct in the Ellenbogen et al. studies [4,8]. As a result, performance was quite high in both the sleep and wake control conditions. This may have resulted in overlearning of the word pairs that could have masked an otherwise more robust sleep-wake difference in word pair recall. Yet in a 2^nd^ similar study using the same protocol with audio/visual word pairs learned prior to a daytime nap, it was shown that interference learning had a similar impact on participants who slept, compared to those who remained awake during the 1-hr training-retest interval [20].

Taken together, the Ellenbogen studies provide some of the best evidence that sleep supports a key aspect of consolidation theory: resistance to interference. However, subsequent studies have not uniformly supported this hypothesis, and a direct replication of Ellenbogen et al.’s original studies has not yet been attempted.

### The Present study

In this well-powered study, we replicate the second study by Ellenbogen et al. (2009). As reported in that study, we expected that interference learning would lead to a sleep-wake difference significantly greater than that observed when no interference learning was undertaken by participants (i.e., a significant sleep/wake × no-interference/interference interaction.

In this study we also examined the impact of intrinsic motivation on memory performance. Past research has shown that providing a monetary reward (extrinsic reward) for performing better on a task (e.g., a simple typing task) led to greater task improvement following sleep compared to wake [21]. However, one study from our lab was unable to replicate this effect using a declarative picture pairs task [22]. More recently, our lab has examined *intrinsic* motivation (the desire to perform a task well for its own sake) as a potential modulator of the sleep-wake differences in memory. One study from our lab showed that intrinsic motivation correlated with better task acquisition *and* consolidation (change in performance from training to retest), but that the sleep/wake effect was not impacted by intrinsic motivation [23]. As in that study, we analyzed four items from the Intrinsic Motivation Inventory [24,25] and their relationship to memory performance following sleep and wake. We hypothesized that greater intrinsic motivation would be associated with better declarative memory at training and better retention across the 12-hr training-retest interval, but that intrinsic motivation would not provide an added memory benefit to those who slept as opposed to stayed awake during the retention interval.

## Methods

### Participants

Participants were 97 university students (age: 23.2±1.9yrs, range: 18-28, 59 female). Participants were asked to sleep well the night before the study and not to exceed their usual daily caffeine intake. Of the 108 participants who completed the study, eleven were excluded from analysis for 1) taking stimulant and/or antidepressant medications (n=9) or 2) reporting that they obtained >2hrs less than their typical total sleep time the night before the study (n=2). All participants provided informed consent to participate in the study. The study was approved by the Prisma Health System IRB and the Furman University IRB.

### Paired Associates Task

Here we used the paired associates task from Ellenbogen et al. (2009) [8], designed to provide a within-subjects manipulation of interference. This represented a methodological improvement over the Ellenbogen et al. (2006) study [4], in which sleep/wake condition and interference condition were treated as between-subjects variables, decreasing the statistical power of the design. Stimuli were randomly selected two-syllable words from the Toronto Word Pool [26], used to create three lists of 20 word pairs (A_1−20_-B_1−20_, A_21−40_-B_21−40_, A_41−60_-B_41−60_) and a fourth list used during the interference learning phase (A_41−60_,C_1−20_) (see Figure 1). Word pairs were “matched for imageability, frequency of use, and concreteness” [8]. Examples of word pairs: A-B pair: CARPET-BUBBLE / A-C pair: CARPET-PAPER (same cue word, different target word). The paired associates task was implemented using OpenSesame stimulus presentation software [27]. Assignment of words to list was counterbalanced using three orders (list 1, 2, 3; list 2, 3, 1; list 3, 1, 2), ensuring that each word list was equally represented in each serial position.

**Figure 1.**
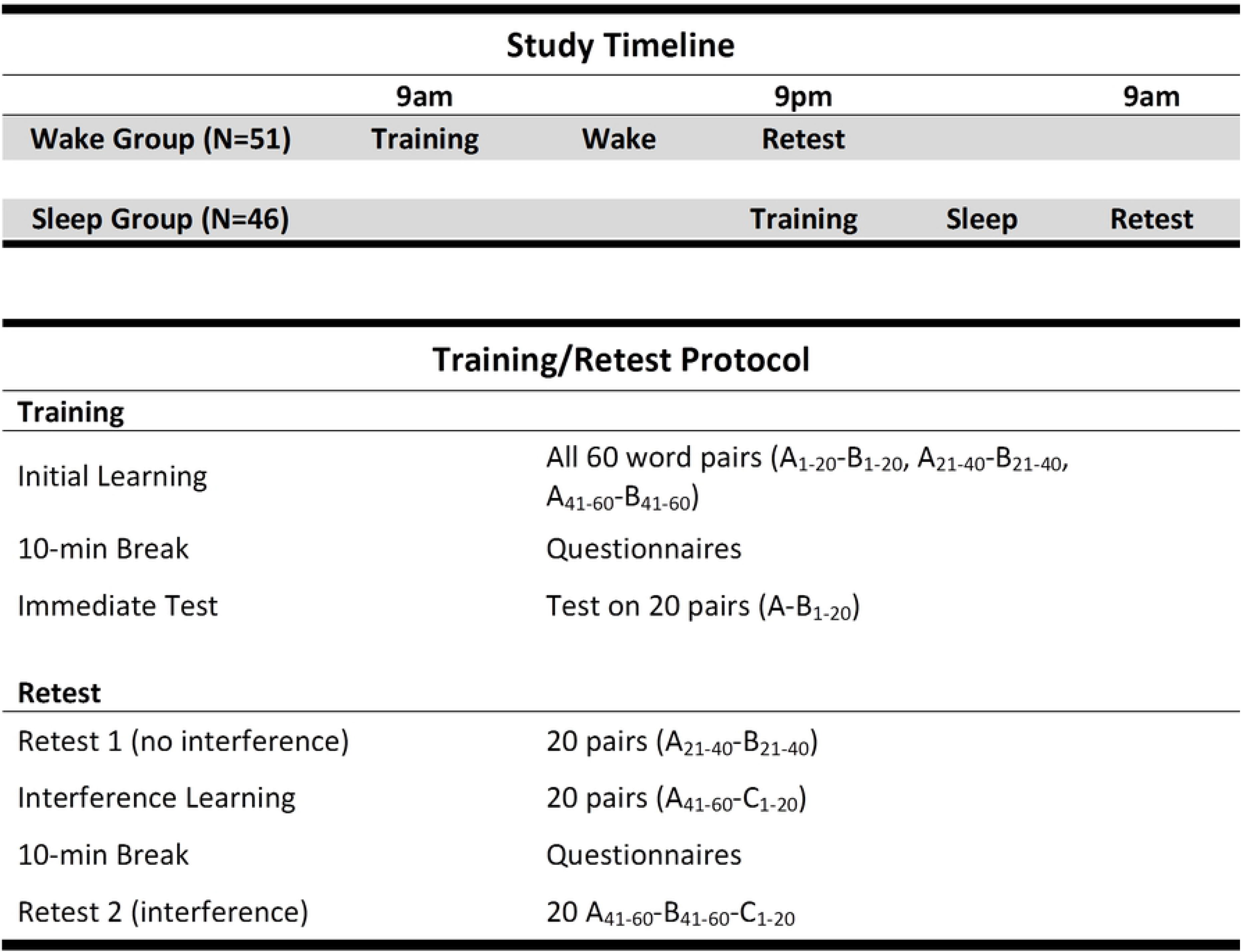
*Study Timeline and Protocol*. The top part shows the timeline for the Sleep and Wake participants from training to retest. The bottom part describes the word pairs training and retest protocol.

During the initial training session in the morning (9am) or evening (9pm), all 60 word pairs were presented for 7s each. Participants were instructed to memorize the pairs but were not given specific strategies about how to memorize them. Following presentation of the word pairs, participants were then quizzed on all 60 pairs, during which they were presented the first word of the pair (A) and were instructed to enter the word that was paired with it (B) in a text box next to the cue word. If participants correctly typed the B word, the pair would not appear again during quizzing. If a response was not correct, the correct pairing would appear on the screen and the item would appear again for another attempt after a random interval. The quizzing phase was complete once all of the A-B pairs were entered correctly one time. There was then a 10-minute break period, during which participants completed paper-pencil study forms. After the 10-minute break, participants were tested on 20 of the 60 word pairs (e.g., A_1−20_-B_1−20_) as a measure of immediate memory (Immediate Test). During this test no feedback was provided after a response was entered.

Twelve hours later, after a night of sleep (9am) or a day of wake (9pm), participants returned for the retest session. During the first part of this session participants were tested on a different set of 20 word pairs from the original set of 60 (e.g, A_21−40_-B_21−40_; Retest 1), a test of memory for word pairs uninfluenced by interference learning. Following this memory test, participants completed the interference learning phase by learning a new set of 20 word pairs that overlapped with the remaining 20 word pairs from the original learning session (e.g., A_41−60_-B_41−60_). This A-C list contained the same A words that were learned originally, paired with 20 new C words (A_41−60_-C_1−20_). The 20 A-C pairs (A_41−60_-C_1−20_) were learned in the same way as the original 60 word pairs, with presentation of the word pairs followed by quizzing with feedback until all pairs were correctly answered one time. There was a 10-minute break following A-C interference learning, during which participants completed the remaining study forms. After the break participants were tested on both the B_41−60_ and C_1−20_ words that had been paired with A_41−60_ (post-interference Retest 2). The A word for each pair was presented and participants attempted to enter both the B word (from the training session) and C word (from the interference session) that had been paired with it.

### Procedure

All sessions were conducted either in a lecture hall at the University of South Carolina School of Medicine Greenville, or in a computer laboratory at nearby Furman University. Participants either participated in a Wake group that completed the training session at 9am and the retest session at 9pm, or a Sleep group that trained at 9pm and retested the next morning at 9am (Figure 1). Upon arrival, participants signed the consent form, and then trained on the 60 word pairs. Following training, during the 10-minute break that preceded an Immediate Test on the A_1−20_-B_1−20_ word pairs, participants completed a set of forms that included a demographics form, a 3-day retrospective sleep log, alertness/sleepiness scales, and the Epworth Sleepiness Scale [28], a measure of the likelihood of falling asleep in eight situations (e.g., sitting and reading a book). The alertness/sleepiness scales consisted of two visual analog scales: “How would you describe your ability to concentrate right now?” and “How refreshed do you feel right now?”, and the Stanford Sleepiness Scale [29], which measures how alert/sleepy the participant feels at the moment. Following the immediate test on the A-B word pairs, participants were free to leave, and were instructed to limit caffeine intake and not to nap (Wake participants) prior to returning for the retest session. The duration of the training session was 30-40min.

When participants returned 12 hours later, they were retested on 20 of the items learned during the training session (e.g, A_21−40_-B_21−40_; Retest 1). This initial test was followed by interference learning of the A-C word pairs. During the 10-minute break that followed A-C interference learning, participants completed forms that included the alertness/sleepiness scales, a sleep or wake log that documented participant activities during the training-retest interval, and a ‘Task Perception Inventory’ that included four items from the from the Intrinsic Motivation Inventory (IMI) [24,25,30]: “I enjoyed doing the word pairs task very much” (Enjoyment), “I didn’t try very hard to do well at the word pairs task” (Effort), “I think I am pretty good at the word pairs task” (Competence), and “I felt very tense while doing the word pairs task” (Tension). These questions were answered on a 5-point Likert scale: “Not at all true” to “Very True”. Participants then answered two questions on a visual analog scale (measured in mm): “How motivated were you to do well on the word pairs task?” (Motivation) and “How much did you think about the word pairs task before the retest session?” (Thinking about). After completing the final test (A_41−60_-B_41−60_-C_1−20_; Retest 2), the study was concluded and participants were paid for their participation. The duration of the retest session was 30−40min.

#### Power Analysis

To ensure the current replication study was well-powered to detect the effects reported in Ellenbogen et al. (2009), effect size was calculated from Ellenbogen et al.’s crucial Sleep x Interference interaction effect, which was the primary statistic demonstrating that the effect of interference on memory depended on sleep/wake condition (partial eta^2^ = 0.099). Using this value, N=76 would be required to achieve power=0.80. The sample used in this replication (N=97), which was more than double the sample size of the original study, yielded statistical power=0.89 to detect the effect of interest.

## Results

### Participant Data

Means±SEMs are presented for the Sleep group first. Participants in the sleep and wake conditions reported similar sleep log and demographic data: Usual bedtime: 10:58pm±7.8min, 11:02pm±8.4min, p=0.72; Usual total sleep time: 7.2±0.1hrs, 7.0±0.1, p=0.21; Total sleep time the night before the study: 7.3±0.2hrs, 7.2±0.2hrs, p=0.53; Epworth Sleepiness Scale score: 8.3±0.6, 7.7±0.6, p=0.47. Self-report measures of sleepiness and alertness revealed that participants in the Wake group reported feeling more refreshed (39.7±3.9mm, 50.8±3.7, p=0.04), marginally more alert (SSS: 3.0±0.2, 3.4±0.2, p=0.09), but not better “able to concentrate” (58.4±3.6mm, 53.2±3.7, p=0.33) than those in the Sleep group. Sleep subjects went to bed earlier than wake subjects the night before the study (11:22pm±8.4min, 12:23±10.2min, p<0.001) but, as noted above, total sleep times were comparable.

### Paired Associates Performance

#### Training

At training, Sleep and Wake groups performed similarly on the immediate test of B words (Percent correct: 82.1±2.7% (16.4 pairs), 83.8±2.6% (16.8), t_(95)_=0.46, p=0.64; Trials to reach criterion: 152.0±9.1, 147.0±14.5, t_(95)_=0.30, p=0.77).

A 2×2 ANOVA was conducted to examine the effects of Sleep v. Wake (between subjects) and Interference v. No-Interference (within subjects) on memory for the B words at retest. As expected, there was a significant main effect of Sleep v. Wake, with memory for B words being superior in the Sleep group (F_(1,95)_=4.44, p=0.038). There was also a significant main effect of Interference v. No-Interference, with memory being superior for No-Interference pairs (F_(1,95)_=70.61, p<.0001). However, crucially, there was no Sleep x Interference interaction effect (F_(1,95)_=0.38, p=0.54), indicating that the effect of sleep on memory did not differ between Interference v. No-Interference pairs (Figure 1). Order of list presentation did not affect recall on the immediate test (at training) (Order 1: 16.1±0.6, Order 2: 16.2±0.8, Order 3: 17.3±0.6, F_(2,94)_=1.10, p=0.34).

#### Retest

Prior to interference training (Retest 1 – no interference)), Sleep participants recalled more B words than Wake participants: 72.7±2.8%, 62.3±3.5%, t_(95)_=2.35, p=0.02. However, the Sleep-Wake difference for post-interference retention (Retest 2 – Interference) did not reach significance: 55.9±3.1%, 47.7±4.1, t_(95)_=1.59, p=0.12. Importantly, there was no interaction between sleep/wake condition and interference condition, with Sleep participants dropping 16.9±2.3% from Retest 1 to Retest 2, and Wake participants dropping 14.6±3.0% (F_(1,95)_=0.38, p=0.54; Figure 2).

**Figure 2.**
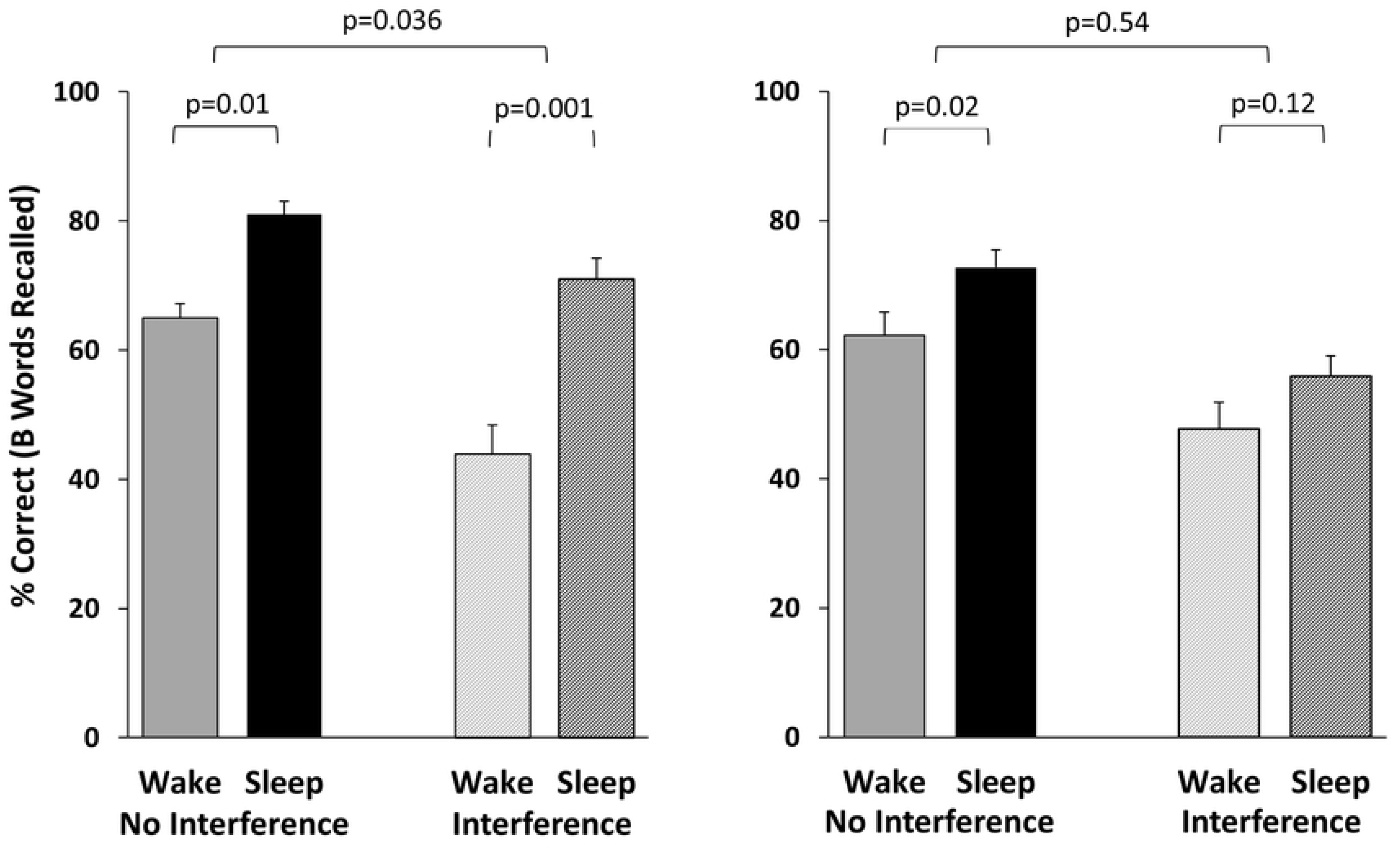
*Paired Associates Results.* Results from Ellenbogen et al. (2009) (Left) are compared to findings from the current study (Right). Results for “no interference” represent recall from the initial retest, prior to interference learning. “Interference” results represent recall of originally-learned B words following interference learning.

#### Interference (A-C) training

Sleep and Wake groups recalled a similar percentage of interference (AC) word pairs: 78.6±3.1, 76.5±3.6, t_(95)_=0.45, p=0.65). The number of trials required to reach criterion for A-C training was also similar between groups (36.6±2.0, 33.9±1.9, t_(95)_=0.97, p=0.34).

### Intrinsic Motivation

Table 2 shows the correlation of memory performance with measures of task perception (items from the Intrinsic Motivation Inventory) and the ‘Motivate’ and ‘Think About’ visual analog scales. We found that three items in particular (Enjoyment, Competence, and Tension) correlated with performance (correct items) at training, Retest 1 (pre-interference learning), and Retest 2 (post-interference learning). The VAS question regarding how motivated participants were to do well on the task also correlated with recall at these three time points. However, there were no significant correlations between any of these questionnaire measures and change in performance from training to retest. Fisher’s Z tests revealed no significant differences between correlations for Sleep vs Wake participants.

## Discussion

The studies by Ellenbogen et al. [4,8]that form the basis for the current replication were important because they demonstrated that sleep does more than passively protect memories from interference. These studies strongly suggested that sleep stabilized memories over the training-retest interval, such that interference learning after a night of sleep had a much smaller impact on memory than after a day of wakefulness. According to the authors, this central finding “demonstrates the active role of sleep in consolidating memory” [8].

In the present study, as in the Ellenbogen studies, we found that, prior to interference training, those who slept had better memory for the word pairs than those who were awake during the 12hr interval. However, we did not replicate the finding that sleep strengthens memories against interference learning. Instead, we observed that the Sleep and Wake groups both showed a significant and very similar retroactive interference effect.

One reason for the discrepancy between our findings and those of Ellenbogen et al. may relate to differences in testing protocol. In their first study [7], participants were required to get each word pair correct twice during training, which resulted in a very high rate of memory performance at retest in the sleep group (with sleep participants getting 94% of items correct at Retest 1). This suggested a ceiling effect that may have artificially attenuated the effect of sleep on recall of non-interference word pairs (Retest 1, where performance levels were highest), thus creating a spurious interaction between sleep-wake condition and interference condition. In their 2009 follow-up study [8], this ceiling effect was no longer apparent -- although some participants were again required to answer each item correctly twice, others were required to answer them correctly only once. In that 2009 study, the sleep-wake x interference-no interference interaction was slightly diminished compared to their original 2006 finding, though still statistically significant. In contrast, in the current study, all participants were required to get each item correct only once, which kept the training protocol the same for all participants, and reduced the risk of ceiling effects at initial retest (Retest 1).

Given the likelihood that participants were over-trained in both the Sheth, et al. (2012) (3x correct at training) and the Ellenbogen et al. (2006) study (2x correct), having participants answer items correctly only one time in the present study should have been sufficient to replicate the interference effect observed in Ellenbogen, et al. (2009), without encountering ceiling effects. Our findings, however, are instead in line with those of Alger, et al. (2012), who examined the impact of interference learning after a 1-hr training-retest interval using an audio/visual word pairs task. They found that a 60-min daytime nap did not reduce the impact of interference learning compared to wake. Thus, it is possible that the apparent ability of sleep to protect against interference in Ellenbogen et al.’s original paper was the spurious result of ceiling effects to which our current data were not susceptible.

The difference between our findings and those of Ellenbogen et al. is not likely due to a lack of statistical power. The sample size of our study was more than 2x as large at that used in Ellenbogen et al. (2009), and we were powered to replicate their effect at 0.90. The observed effect size for our sleep/wake x no-interference/interference interaction was quite small (partial eta-squared = 0.003), and the Ellenbogen et al. (2009) effect (partial eta-squared = 0.099) fell outside the 90% confidence interval of our effect (Figure 3). Importantly, although the 90% confidence interval on our effect is consistent with a small protective effect of sleep against interference, this hypothetical effect size would have been too small for Ellenbogen’s study to meaningfully detect.

**Figure 3.**
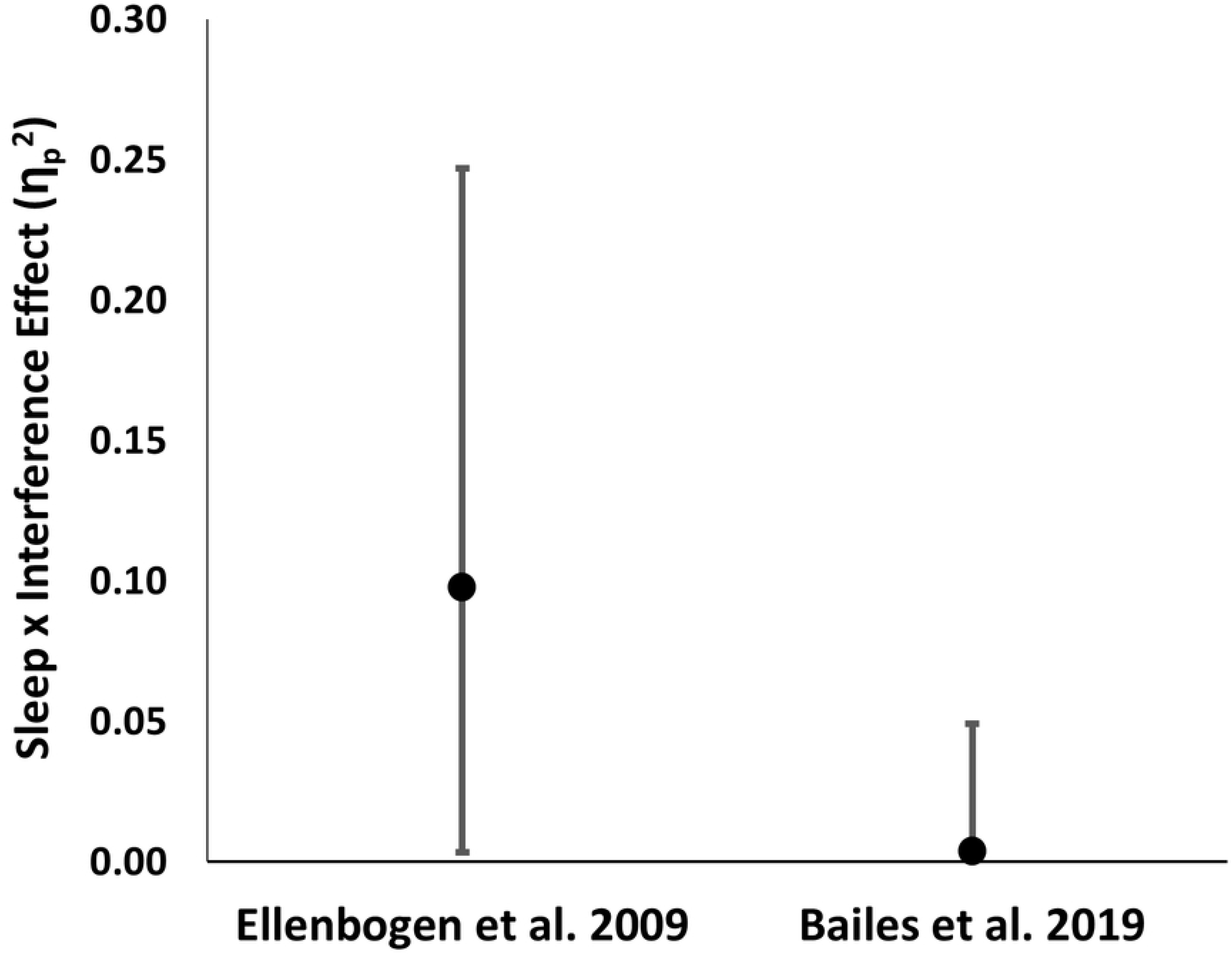
*Effect Size Comparison.* Points represent the size of the Sleep × Interference interaction effect as reported by Ellenbogen et al. (2009), and in the current study (partial eta squared). Error bars represent the 90% confidence interval on this effect. The Ellenbogen et al. 2009 effect falls well outside the 90% confidence interval of the current, more precise effect size estimate.

A second aim of this study was to examine intrinsic motivation and its relationship to measures of memory performance. A previous study from our lab found that providing participants an extrinsic reward for performance ($1 per correct answer) resulted in better recall of visual paired associates (picture pairs) compared to no reward, but that this reward effect was not different between sleep and wake participants [22]. We have also shown that increased intrinsic motivation (the desire to do well even in the absence of extrinsic rewards), measured by responses to items from the Intrinsic Motivation Inventory [24] used in the present study correlated with better acquisition of visual paired associates (picture-object pairs) at training, and also with change in performance from training to retest (retention) [23]. In the present study, we observed a similar positive correlation between intrinsic motivation and training performance. However, motivation did not correlate with memory consolidation (change in performance from training to retest), nor did intrinsic motivation favor performance in the sleep (v. wake) participants. These findings point to the potential importance of intrinsic motivation for optimizing learning and memory, but that intrinsic motivation may not be an important modulator of sleep-wake differences in memory.

This replication study adds to our understanding of the functional role of sleep in memory processing. While studies by Ellenbogen et al. (2006, 2009) seemed to provide strong evidence that sleep is actively stabilizing memories against interference, our findings draw this now commonly accepted finding into question. This study also highlights the importance of independent replication of seminal findings in the field, and suggests that challenges remain as we continue to define the specific nature of sleep’s role in memory processing.

**Table 1.** Correlations between indices of intrinsic motivation from the Intrinsic Motivation Inventory, training performance and change in performance from training to retest.

|  |  | Training<br>(Immediate test) |  | Retest1<br>(No interference) |  | Retest2<br>(Interference) |  |
| --- | --- | --- | --- | --- | --- | --- | --- |
|  |  | r | p | r | p | r | p |
| Intrinsic<br>Motivation<br>Inventory | Enjoyment | .42 | **<.001** | .42 | **<.001** | .36 | **<.001** |
|  | Effort | .14 | .17 | .14 | .17 | .05 | .63 |
|  | Competence | .55 | **<.001** | .49 | **<.001** | .39 | **<.001** |
|  | Tension | -.40 | **<.001** | -.26 | .01 | -.19 | .06 |
|  | Total Score | .45 | **<.001** | .39 | **<.001** | .38 | **<.001** |
| Visual<br>Analog<br>Scales | Motivation | .31 | **.002** | .23 | .02 | .23 | .03 |
|  | Thinking<br>About | -.05 | .65 | .14 | .17 | .10 | .35 |

Note: Retest1: First test of the retest session, prior to interference learning ( $A_{21-40}$ - $B_{21-40}$ ): Retest2: Post-interference learning retest on the original B words ( $A_{41-60}$ - $B_{41-60}$ ). Bold: significant correlations.

## Acknowledgements

We thank Ted Summer for his assistance in programming the OpenSesame version of the paired associates task, and Daley Rhines for assistance with data collection. This research was funded in part by grant 1R15MH107891 from NIMH.

